# Cerebral blood flow in elastin haploinsufficient and 3xTg-AD mice

**DOI:** 10.1101/2025.08.18.670895

**Authors:** Abigail E. Cullen, Emily H. Reeve, Nick R. Winder, Grant D. Henson, Nayantara Arora, Thomas Leonhardt, Ainsley Hogan, Sahana Krishna Kumaran, Naly Setthavonsack, Victoria Krajbich, Nabil J. Alkayed, Martin M. Pike, Randall L. Woltjer, Ashley E. Walker

**Affiliations:** Human Physiology, University of Oregon, Eugene, OR, USA; Pathology and Laboratory Medicine, School of Medicine, Oregon Health and Science University, Portland, OR, USA; Knight Cardiovascular Institute, Oregon Health and Science University, Portland, OR, USA; Biomedical Engineering, Advanced Imaging Research Center, Oregon Health and Science University, Portland, OR, USA

**Keywords:** cerebrovascular reactivity, elastin haploinsufficiency, neuroinflammation, Alzheimer’s disease

## Abstract

Artery structural properties and Alzheimer’s disease (AD) pathology are individually associated with impaired cerebrovascular function; however, the interaction of these factors is unclear. Furthermore, while elastin haploinsufficient (*Eln*^*+/−*^) mice are known to have impaired cerebrovascular function, sex differences for this effect have not been previously studied. To answer these questions, we crossed middle-aged and old *Eln*^*+/−*^mice with 3xTg-AD mice. We measured cerebral blood flow (CBF) using arterial spin labeling MRI at rest and during hypercapnia to calculate cerebrovascular reactivity (CVR). We also assessed neuroinflammation by microglia and astrocyte cell counts. We found that *Eln*^*+/−*^ mice had lower resting blood flow rate in the cerebral cortex compared with *Eln*^*+/+*^ mice, but *Eln*^*+/−*^ mice had an intact hypercapnic response, resulting in better CVR compared with *Eln*^*+/+*^ in hippocampus. Sex did not impact resting blood flow or CVR. 3xTg-AD mice had a lower resting CBF than non-AD mice, and there was an interaction between *Eln* genotype and AD mutations on CVR, such that *Eln*^*+/−*^x 3xTg-AD mice had the poorest hippocampal CVR of all groups. Glia cell counts were highly dependent on brain region, with *Eln*^*+/−*^ having more microglia but fewer astrocytes, while 3xTg-AD having higher both microglia and astrocytes. While sex also impacted glial cell counts, we found no interactions between sex and *Eln* genotype. Our results demonstrate that elastin haploinsufficiency and AD mutations individually result in lower resting CBF, and the combination of these leads to impaired CVR.

**NEW & NOTEWORTHY:** The findings of this study demonstrate that elastin haploinsufficiency leads to lower resting cerebral blood flow, but also greater cerebrovascular reactivity. However, elastin haploinsufficiency interacts with Alzheimer’s disease mutations to impair cerebrovascular reactivity. These results suggest that multiple insults, such as changes to the extracellular matrix combined with genetic risk factors, are needed to impact cerebrovascular reactivity.

## INTRODUCTION

Elastin is a primary component of the vascular extracellular matrix that contributes to the elasticity of arteries and impacts cerebrovascular function (1). Elastin haploinsufficient mice (*Eln*^*+/−*^) have stiffer arteries, at least for the large elastic arteries, compared with wild-type mice (2, 3). These differences in artery structure coincide with impairments to cerebrovascular function. Our previous work demonstrates that *Eln*^*+/−*^ mice have impaired cerebral artery endothelium-dependent vasodilation and a greater vasoconstrictor responsiveness to angiotensin II (2, 4). *Eln*^*+/−*^ mice also have a lower cerebral blood flow (CBF) in the cerebral cortex than wild-type mice (5). However, resting CBF provides limited information about cerebrovascular health, while cerebrovascular reactivity (CVR), such as the response to a hypercapnic stimulus, is more indicative of dynamic cerebrovascular function. Moreover, resting CBF and cerebrovascular reactivity (CVR) do not always follow the same trends. For example, in human subjects, greater aortic stiffness is associated with lower basal CBF, but higher CVR (6). Thus, measuring CVR provides a better understanding of the effects of elastin haploinsufficiency on cerebrovascular health.

There are significant sex differences in the trajectories of changes in arterial stiffness and cerebrovascular function with advancing age. The rate of increase in carotid artery stiffness is faster with aging in females compared with age-matched males (7). In addition, the age-related declines in CBF and CVR are greater in females than males (8, 9). Sex differences are also found in aged *Eln*^*+/−*^ mice, with the trajectories for changes in structural properties of the large arteries being different between male and female *Eln*^*+/−*^ mice (10). However, sex differences in cerebrovascular function in *Eln*^*+/−*^ have not been previously studied.

An important consequence of poor cerebrovascular health is an increased risk of dementia and worsening of Alzheimer’s disease (AD). AD is associated with lower CVR (11-14), which may be due to the vascular effects of AD pathology, such as increased amyloid-β (Aβ) and hyperphosphorylated tau. Aβ causes vasoconstriction in the cerebral vasculature (15), and mice with increased Aβ production have lower CBF (16). In addition, mutant tau overexpression is associated with altered cerebral blood vessel structure and lower CBF in mice (17). However, it is unclear whether changes to the extracellular matrix impact the effects of AD pathology on the cerebral vasculature. To understand this interaction, we crossed *Eln*^*+/−*^ mice with the 3xTg model of AD, which has AD-related mutations that lead to Aβ accumulation and tau hyperphosphorylation.

Neuroinflammation can impair cerebrovascular function in AD, while reduced CBF can lead to hypoxia-induced neuroinflammation (18). Neuroinflammation is characterized by an increase in glial cell activation, notably astrocytes and microglia (19). Chronically high levels of neuroinflammation can contribute to cognitive impairment, blood-brain barrier disruption, and increased Aβ production(19). There are sex-specific differences in neuroinflammation, specifically microglia isolated from female brains are more pro-inflammatory than those from male brains (20), and the microglia from female brains are more strongly related to AD pathogenesis (21). Yet, the impact of elastin haploinsufficiency on neuroinflammation has not been previously studied.

In these studies, we set out to test two hypotheses: 1) sex and elastin haploinsufficiency interact to impact cerebrovascular function and neuroinflammation, such that female *Eln*^*+/−*^ mice would have worse CVR and neuroinflammatory markers compared with male *Eln*^*+/−*^ mice, and *Eln*^*+/****+***^ of either sex; and 2) AD pathology and elastin haploinsufficiency interact to impact cerebrovascular function and neuroinflammation, such that *Eln*^*+/−*^ x 3xTg-AD mice would have worse cerebrovascular reactivity and neuroinflammatory markers compared with *Eln*^*+/****+***^ x 3xTg-AD mice, and *Eln*^*+/****+***^ or *Eln*^*+/−*^ mice without the 3xTg-AD variants.

## MATERIALS AND METHODS

### Animals

*Eln*^*+/−*^ mice were transferred from the University of Utah and the National Institutes of Health(5) were rederived with pathogen-free C57BL/6J at the University of Oregon. The 3xTg-AD mice were originally created by Frank LaFerla at the University of California, Irvine and were obtained from the Jackson Laboratory (#004807). We bred *Eln*^*+/−*^ mice with the 3xTg-AD mice at the University of Oregon and studied the F2 generation. Two months before magnetic resonance imaging (MRI), mice were transferred to Oregon Health and Science University. We studied males and females at ages of 14 to 22 mo. The age range was due to research delays associated with the COVID-19 pandemic. However, there were no differences in mean age between groups (*Eln*^*+/+*^: 16.2 mo, *Eln*^*+/−*^: 16.4 mo, *Eln*^*+/+*^x 3xTg-AD: 16.8 mo, *Eln*^*+/−*^x 3xTg-AD: 15.1 mo; 16.2±2.6 mo, p=0.472). Further, there were no differences between male and female age in any groups (M *Eln*^*+/+*^: 16.2 mo, F *Eln*^*+/+*^: 16.2 mo, M *Eln*^*+/−*^: 16.6 mo, F *Eln*^*+/−*^: 16.3 mo, M *Eln*^*+/+*^x 3xTg-AD: 16.1 mo, F *Eln*^*+/+*^x 3xTg-AD: 18.5 mo, M *Eln*^*+/−*^x 3xTg-AD: 14.6 mo; F *Eln*^*+/−*^x 3xTg-AD: 15.9 mo). After the MRI, mice were sacrificed by cardiac puncture and perfused with saline. Following saline, 4% paraformaldehyde was perfused to preserve the brain for histology. All mice were housed in an animal care facility on a 12/12 h light-dark cycle at 24°C with ad libitum access to standard chow (Lab Diet, Picolab Rodent Diet 20-5053, University of Oregon; Lab Diet, Laboratory Rodent Diet-5001, Oregon Health and Science University) and water. All animal procedures conformed to the *Guide to the Care and Use of Laboratory Animals* and were approved by the Institutional Animal Care and Use Committee at the University of Oregon and Oregon Health and Science University.

### Arterial Spin Labeled MRI

MRI was performed at the OHSU Advanced Imaging Research Center using a Bruker-Biospin 11.75 T small animal MR system with a ParaVision 6.0 software platform, 10 cm inner diameter gradient set with a 72 mm (ID) and 60 mm (length) RF resonator for transmitting and an actively decoupled mouse head surface coil for receiving. Mice were anesthetized with a ketamine/xylazine mixture (1.0 mg xylazine/7 mg ketamine/100 g) in combination with low isoflurane (0.75%) in 100% oxygen. The mice were positioned with heads immobilized on an animal cradle. Body temperature of the mice was monitored and maintained at 37^◦^C while monitoring respiration (SA Instruments, Stony Brook, NY, United States). For each mouse, a coronal 25-slice T2-weighted image was acquired (ParaVision spin echo RARE, 256 × 256 matrix, 125 µm in-plane resolution, 0.5 mm slice width, TR 4000 ms, TE effective 23.64 ms, RARE factor 8, 2 averages). These T2-weighted anatomical scans were used for positioning the blood flow image slice at a consistent position approximately 1.75 mm anterior to the anterior commissure. CBF (ml/min/100 g) was measured using ASL, employing the flow-sensitive alternating inversion recovery rapid acquisition with relaxation enhancement pulse sequence (ParaVision FAIR-RARE), with TE/TR = 45.2/10000 ms, slice thickness =1 mm, number of slices = 1, matrix = 128 × 128, 250 µm in-plane resolution RARE factor = 72, and 23 turbo inversion recovery values ranging from 40 to 4400 ms, and acquisition time of 15 min. This sequence labels the inflowing blood by global inversion of the equilibrium magnetization (22). The ASL sequence was implemented first at the baseline condition with 100% oxygen, and subsequently 10 min after switching to the hypercapnia condition with 95/5% oxygen/carbon dioxide to assess CVR, defined as the percent increase in blood flow with hypercapnia. CBF maps (ml/100 g-min) were generated using the Bruker Paravision ASL perfusion processing macro and exported into JIM 9 software (Xinapse Systems LTD, Northants, United Kingdom) for further processing. Outlier value brain pixels (outside 2 SDs) representing large arteries with high, pulsatile flow were excluded, thus arriving at flows which consistently represent tissue microvascular blood flow. The identical FOV geometry offsets of the T2 and ASL images enabled the ROIs drawn on the T2 image to be readily overlaid onto the corresponding blood flow map for quantification. Mean blood flow was quantified for the whole brain, cortex, hippocampus, and thalamus, each defined anatomically using the corresponding T2 images. ASL map voxels containing ventricular contributions were excluded. The Allen Brain Atlas as a guide to identify ventricles, hippocampus, cortex, and thalamus.

### Immunohistochemistry

After paraformaldehyde perfusion, brains were incubated in 4% paraformaldehyde for 48 h before being embedded in paraffin. Paraffinized brains were cut at 5 um thickness and incubated with primary antibodies for Iba1 (ProteinTech, 10904-1-AP, 1:5000), GFAP (ProteinTech, 16852-1-AP, 1:500), Aβ (4G8, Biolegend, 800709, 1:5000), or tau (PHF1, kind gift from Peter Davis laboratory, 1:5000), followed by development using Vector ELITE ABC kits (Vector Labs, PK-6102). Visualization was done with 3,3’-diaminobenzidine (23). Regions of interest (hippocampus, entorhinal cortex, and thalamus) were determined using the Allen Brain Atlas and images were collected at a consistent area size (600×450 µm) using a Zeiss Axio Imager AZ10 and analyzed using Image J software (NIH, Bethesda, MD, USA) by percent positive area and particle (cell) size.

### Statistical Analyses

Statistical analyses were performed with GraphPad Prism 10.2.3. Two-way analysis of variance (ANOVA) was used to determine interactions between elastin genotype and sex on CBF and immunohistochemistry. Due to small sample sizes, sexes were combined for the comparisons of AD genotypes, resulting in a two-way ANOVA to determine interactions between elastin haploinsufficiency and AD genotype. Significance was set at p < 0.05 and all values shown are means ± standard deviation. Outliers were determined to be values with Z-scores outside the range of −2.0 to 2.0 and were removed from the data set. In cases of a significant F value, post hoc analyses were performed using the Tukey correction for preplanned comparisons. If data did not fit a normal distribution, we used transformation by log(y) or square root(y) to create normal distributions and proceeded with ANOVA and post hoc analysis.

## RESULTS

### Elastin haploinsufficiency has a stronger impact on resting CBF and CVR than sex

Resting CBF in the cortex was significantly lower in *Eln*^*+/−*^ mice compared with *Eln*^*+/****+***^ mice (Figure 1. B, p=0.01). Post hoc analysis indicated a significant difference between the male *Eln*^*+/****+***^ and *Eln*^*+/−*^ mice (Figure 1. B, p=0.02). We found non-significant trends for main effects of elastin haploinsufficiency in the whole brain, hippocampus, and thalamus (Figure 1. A,C,D, all p<0.10). In contrast, sex had no effect on resting CBF of any brain region (Figure 1. A-D, all p>0.05). During hypercapnia, there were no effects of sex or elastin haploinsufficiency on CBF of any brain region (Figure 1. F-I, all p>0.05). We then calculated CVR as the percent change in CBF between resting and hypercapnic CBF. We found a main effect of elastin haploinsufficiency on CVR in the whole brain and hippocampus (Figure 1. K,M, p<0.05) and a trend in the cortex (Figure 1. L, p=0.05), such that the *Eln*^*+/−*^ mice had a greater CVR compared with the *Eln*^*+/****+***^ mice. We found no effect of sex on CVR in any brain region (Figure 1. K-N, p>0.05).

**Figure 1.**
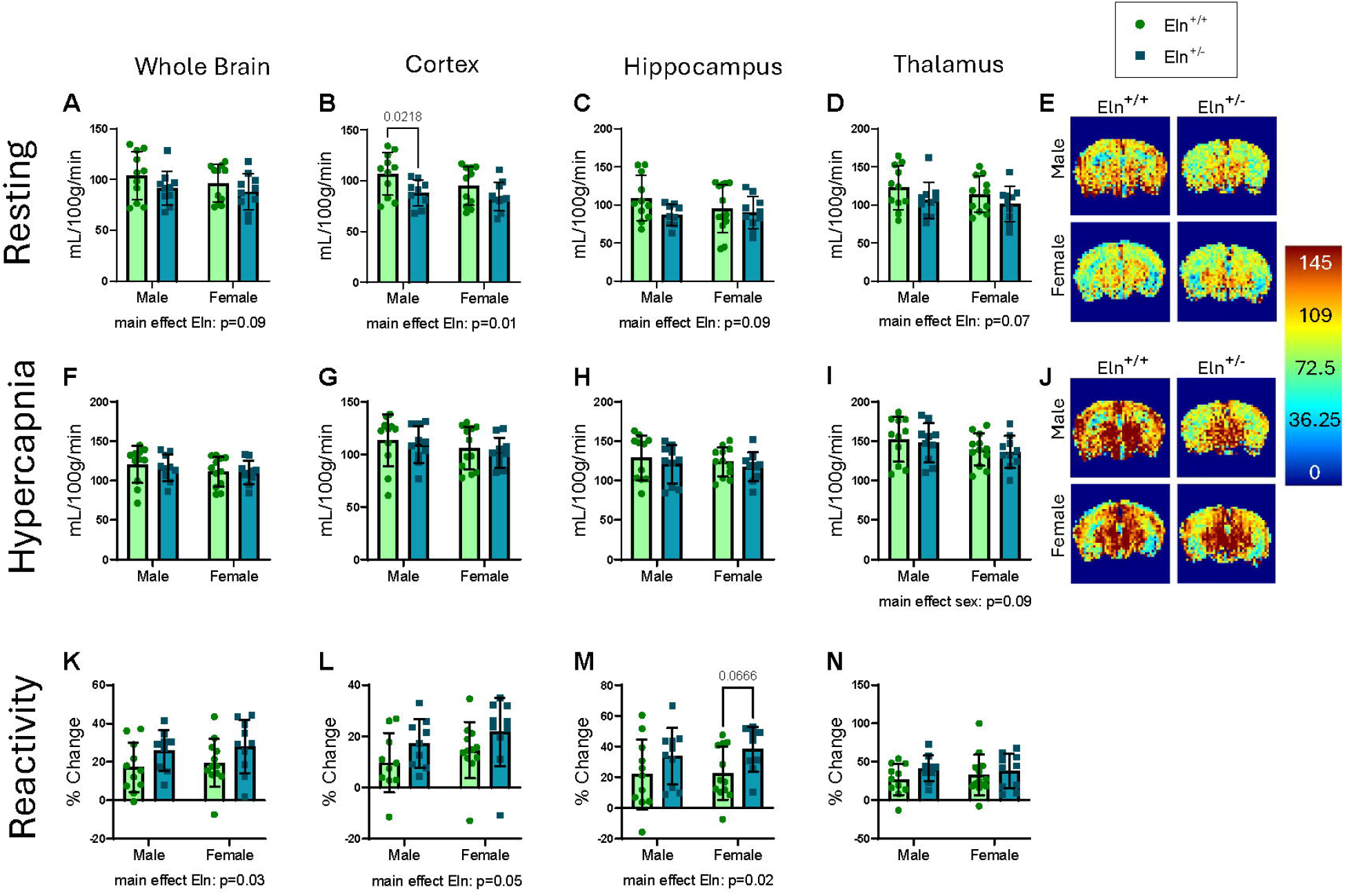
Elastin haploinsufficiency is associated with lower resting CBF and greater CVR. For male and female elastin wild-type (Eln WT)(n=7-13 M/F) and haploinsufficient (HET)(n=7-13 M/F) aged mice (16.2±2.6 mo), blood flow of the whole brain, cortex, hippocampus, and thalamus was measured via arterial spin labeling MRI in resting (A-D) and hypercapnic conditions (F-I), and cerebrovascular reactivity (CVR) was calculated (K-N). Representative images are shown for the resting and hypercapnic conditions in E and J, respectively. All data are shown as mean ± SD. A 2-way ANOVA result p<0.05 was considered significant.

### Sex and elastin haploinsufficiency impact markers of neuroinflammation

We assessed markers of neuroinflammation by measuring Iba1-positive cells, a marker of microglia, and GFAP-positive cells, a marker of activated astrocytes. For Iba1, there was a strong main effect of elastin haploinsufficiency within the entorhinal cortex, indicating more Iba1-positive cells in *Eln*^*+/−*^ mice compared with *Eln*^*+/****+***^ mice (p=0.002), but this was not significantly different in the hippocampus or thalamus (p>0.05, Figure 2. A-C). We also found a main effect of sex on Iba1-positive cells in the entorhinal cortex and thalamus regions (Figure 2. A,C, p<0.05), such that females had a higher cell count in the entorhinal cortex while males had a higher cell count in the thalamus. Post hoc analysis of the entorhinal cortex found significant differences in Iba1 cell count between male *Eln*^*+/****+***^ and *Eln*^*+/−*^ mice, female *Eln*^*+/****+***^ and *Eln*^*+/−*^ mice, and between male and female *Eln*^*+/****+***^ mice. Post hoc analysis of the thalamus data indicated significantly fewer Iba1-positive cells between male *Eln*^*+/****+***^ and female *Eln*^*+/****+***^ mice. When examining the size of the Iba1-positive cells, a marker of cell activation, we found main effects of sex in the entorhinal cortex and hippocampus (Figure 2. D,E, p<0.05), such that females had larger Iba1-positive cells than males. Post hoc analysis showed larger Iba1-positive cells in female *Eln*^*+/−*^ mice compared to male *Eln*^*+/−*^ mice in the hippocampus (Figure 2. E, p=0.05) and a trend for larger cell size in female compared with male Eln+/+ mice in the entorhinal cortex and hippocampus (Figure 2. D,E, p=0.08). For GFAP, we found a main effect of genotype on cell count in the hippocampus (Figure 2. K, p=0.004), such that there were fewer GFAP-positive cells in the *Eln*^*+/−*^ mice compared with *Eln*^*+/****+***^ mice. However, there was no effect of genotype on GFAP-positive cell count in the entorhinal cortex or thalamus (Figure 2. J,L, p>0.05). There were no significant main effects of sex or interactions with sex for GFAP in any brain region (Figure 2. J-L, p>0.05).

**Figure 2.**
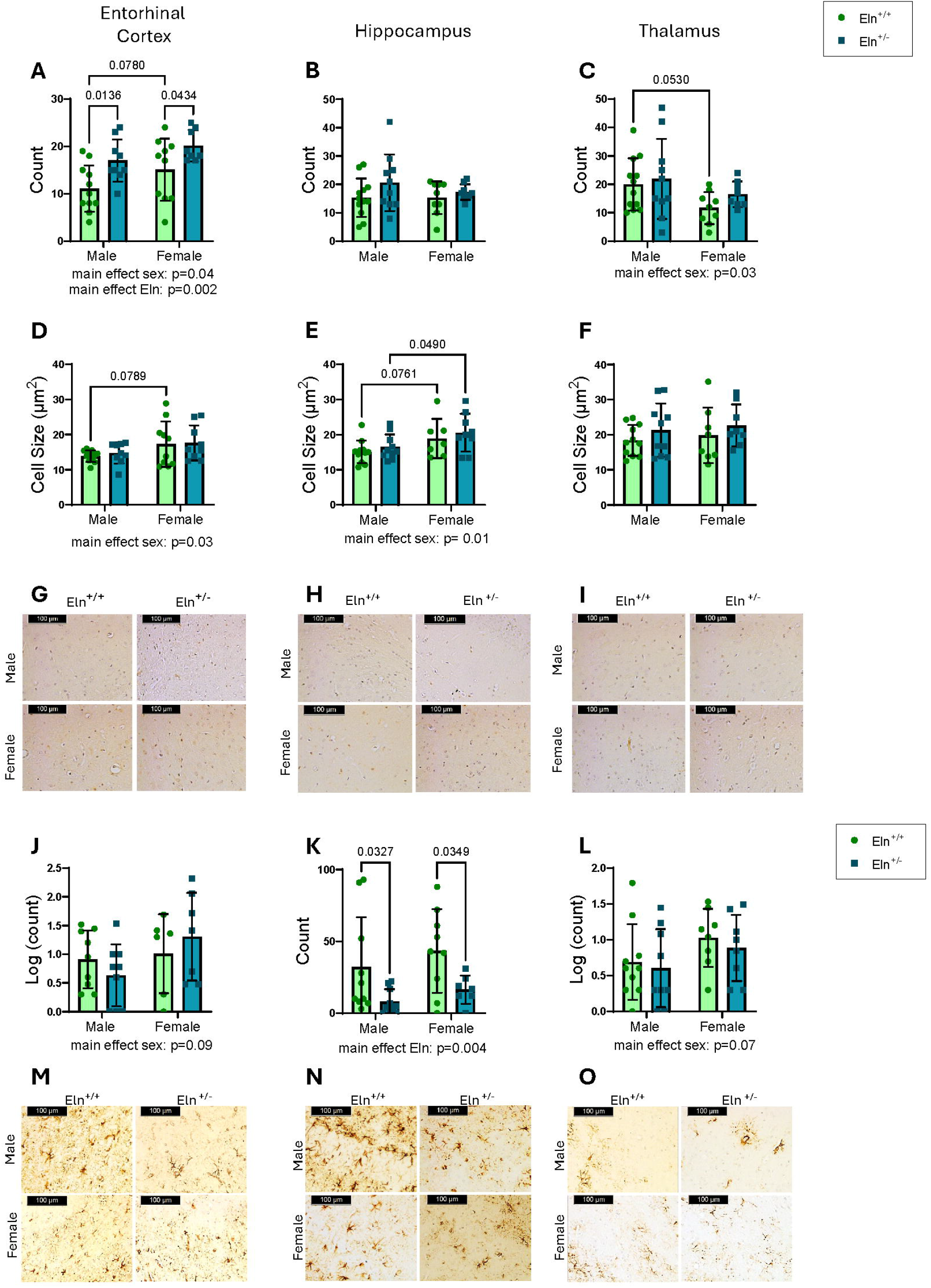
Sex influences microglial, and elastin haploinsufficiency influences microglial and astrocytes. Iba1-positive cell count (A-C) and cell size (D-F) in the entorhinal cortex, hippocampus, and thalamus, respectively, in male and female elastin wild-type (Eln WT)(n=7-13 M/F) and haploinsufficient (HET)(n=7-13 M/F) mice. Representative images are shown in G-I. GFAP-positive cell count for the entorhinal cortex (J), hippocampus (K), and thalamus (L). Representative images are shown in M-O. Non-gaussian data were normalized via log(count). All data are shown as mean ± SD. A 2-way ANOVA result p<0.05 was considered significant.

### CBF is lower in AD mice and CVR is impacted by an interaction of AD and elastin haploinsufficiency

To identify the potential interactions of elastin haploinsufficiency and AD, we crossed *Eln*^*+/−*^ mice with 3xTg-AD mice. CBF in the whole brain, cortex, and thalamus regions was lower in AD mice (Figure 3. A,B,D, p<0.05), and there was an interaction between AD and *Eln* genotype in the hippocampus (Figure 3. C, p=0.03). Post hoc analyses indicated significantly lower CBF in AD x *Eln*^*+/****+***^ mice compared with non-AD *Eln*^*+/****+***^ mice, and lower CBF in non-AD *Eln*^*+/−*^ vs. non-AD *Eln*^*+/****+***^ mice in most brain regions. The lower CBF in AD mice was also found during the hypercapnia condition in all brain regions (Figure 3. F-I, p<0.05). For CVR, there was a significant interaction of AD and elastin haploinsufficiency in the hippocampus (Figure 3. M, p=0.04). Specifically, in non-AD mice, CVR was higher in *Eln*^*+/−*^ compared with *Eln*^*+/****+***^ mice. However, this pattern was reversed in the AD mice, with the AD x *Eln*^*+/****+***^ having higher CVR than the AD x *Eln*^*+/−*^ mice.

**Figure 3.**
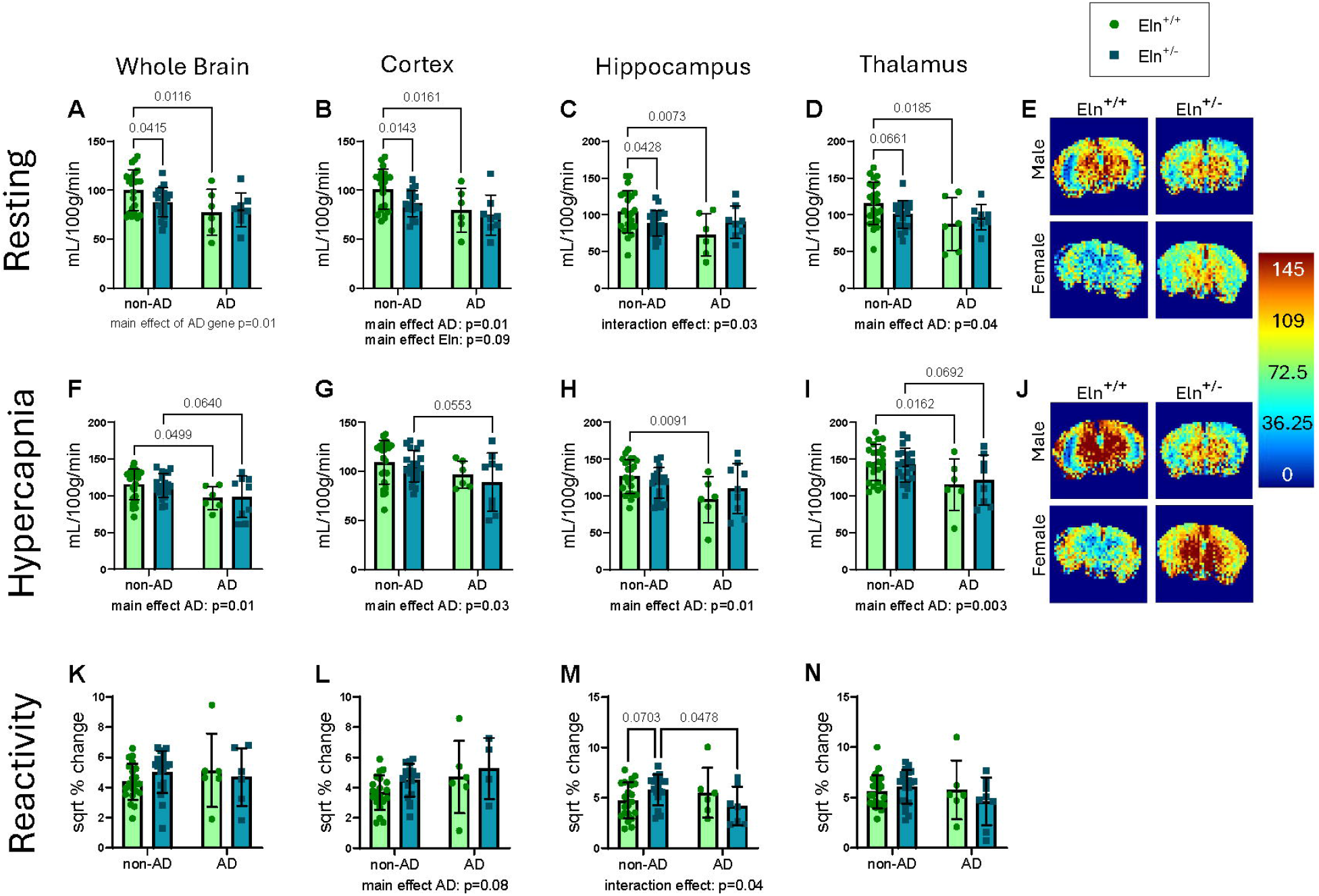
AD mutations significantly impair CBF and interact with elastin haploinsufficiency to dampen CVR. For elastin wild-type (Eln WT)(n=5-6 M/F) and haploinsufficient (HET)(n=5-12 M/F) mice with (AD) and without (non-AD) cross with 3xTg-AD, blood flow of the whole brain, cortex, hippocampus, and thalamus was measured via arterial spin labeling MRI in resting (A-D) and hypercapnic conditions (F-I), and cerebrovascular reactivity (CVR) was calculated (K-N), if data were abnormal a square root (sqrt) transformation was applied to all regions of interest to obtain normality. Representative images are shown for the resting and hypercapnic conditions in E and J, respectively. All data are shown as mean ± SD. A 2-way ANOVA result p<0.05 was considered significant.

### The impacts of AD and elastin haploinsufficiency on neuroinflammation vary across brain regions

a main effect of *Eln* genotype, we found that AD impacted the count of positive Iba1-cells (Figure 4. C, p=0.005), where the AD mice had significantly more Iba1-positive cells compared with the non-AD mice. For Iba1 cell size, we found that in the entorhinal cortex, there was a main effect of *Eln* genotype (Figure 4. D, p=0.01), indicating that the *Eln*^*+/−*^ groups had significantly larger Iba1-positive cells compared to the *Eln*^*+/****+***^ groups. In the hippocampus, we found an interaction effect between the two genotypes (Figure 4. E, p=0.003), resulting in the same pattern as Iba1-positive cell count in the entorhinal cortex. For GFAP, we found a significant main effect of AD genotype on GFAP-positive cell count in all three regions of interest (Figure 4. J-L, all p<0.0001). Post hoc analyses showed AD mice had higher GFAP-positive cell counts than non-AD mice within the *Eln* genotypes for all brain regions (Figure A-C, all p<0.001). We did not find substantial evidence of Aβ plaques or tau tangles in any brain sections.

**Figure 4.**
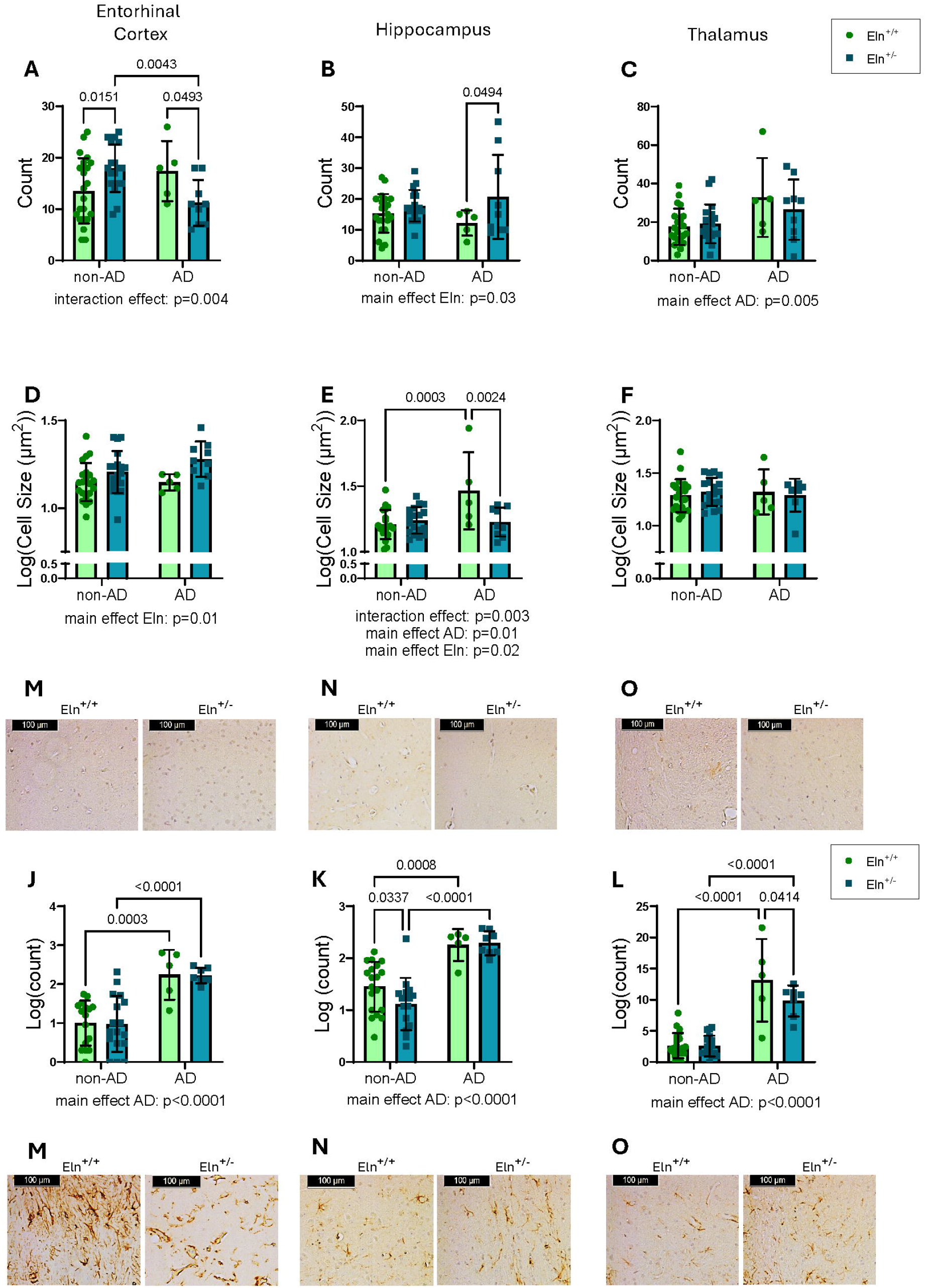
Elastin haploinsufficiency and AD mutations interact to influence microglia and astrocytes. Iba1-positive cell count (A-C) and cell size (D-F) were assessed in the entorhinal cortex, hippocampus, and thalamus, respectively, for elastin wild-type (Eln WT)(n=5-6 M/F) and haploinsufficient (Eln HET)(n=5-12 M/F) mice with (AD) and without (non-AD) cross with 3xTg-AD. GFAP-positive cell count was assessed in the entorhinal cortex (J), hippocampus (K), and thalamus (L)for Eln WT Eln HET mice with (AD) and without (non-AD) cross with 3xTg-AD. Representative images are shown in M-O. Cell count was normalized via log(count). All data are shown as mean ± SD. A 2-way ANOVA result p<0.05 was considered significant.

## DISCUSSION

In this study, we examined the impact of elastin haploinsufficiency on resting CBF, cerebrovascular reactivity, and neuroinflammation, and further explored the interactions of sex and AD-related mutations on these outcomes. We found that although elastin haploinsufficiency leads to lower resting CBF in the cerebral cortex, it is associated with intact hypercapnic response and better CVR in non-AD mice. AD-related mutations were associated with lower resting CBF. Unlike the non-AD mice, the elastin haploinsufficient AD mice did not overcome their lower resting CBF during hypercapnia and had the poorest hippocampal CVR of all groups. This is indicative of an inability to raise CBF to match metabolic demands, potentially predisposing to neuroinflammation and hippocampus-dependent cognitive decline. We found that neuroinflammation was highly dependent on brain region, and in general, female sex was associated with a greater number of microglia. Elastin haploinsufficiency was associated with more microglia but fewer astrocytes, while AD-related mutations were associated with higher number of both activated microglia and astrocytes. In summary, we find that elastin haploinsufficiency and AD-related mutations have disparate impacts on cerebrovascular function and neuroinflammation, which differ by brain region.

Our data, together with previous studies, indicate that elastin haploinsufficiency is associated with lower cortical blood flow throughout the lifespan. Our findings in middle-aged and old mice align with a previous study by Knutsen et al. in young (4 mo) *Eln*^*+/−*^ mice. They found that resting CBF was lower in *Eln*^*+/−*^ mice compared with *Eln*^*+/****+***^ mice (5). They also found a similar pattern for regional impacts, with the most significant effects of elastin haploinsufficiency in the cortex (5). Cortical atrophy is significantly related to cognitive decline in AD (24) and the entorhinal cortex is a crucial site of initiation for AD pathology (25). Thus, this reduction in cortical blood flow with elastin haploinsufficiency could impact cognitive function and AD onset. However, while resting CBF has been associated with cognitive decline and disease(26), a more physiologically relevant measure of dynamic changes in CBF is CVR. Interestingly, we found that elastin haploinsufficiency was associated with higher CVR across multiple brain regions, likely attributed to lower baseline CBF and intact hypercapnic response. Our CVR data agree with Jefferson et al., who found that higher aortic stiffness is associated with higher CVR in cognitively normal older adults. Similar to our study, these correlations with CVR were found across multiple brain regions (6). This greater CVR in elastin haploinsufficient mice and older adults with stiffer large arteries could be mediated by the lower resting CBF, thus representing a ceiling effect in the wild-type and lower stiffness groups. Therefore, it remains unclear whether the higher CVR represents a beneficial or detrimental physiological response.

Beyond the impacts of elastin haploinsufficiency alone, we also find impacts of AD mutations on cerebrovascular function. The presence of AD mutations was associated with significantly lower CBF, but no differences in CVR. In contrast, most studies in humans find that CVR is impaired in older adults with AD, mild cognitive impairment, or cerebral amyloid angiopathy (11, 12, 27). Our negative findings for CVR in the AD mutant mice may be related to our finding of no Aβ plaques in the brains of our mice. Currently, the literature does not have a definitive reason for why CVR is lower in AD, but the vascular dysfunction caused by Aβ plaques is one hypothesis (13). Although we did not find Aβ plaques, the APP and PSEN mutations in these mice likely increased soluble Aβ, which may explain the lower resting CBF. Moreover, our findings align with Shin et al., who found that Aβ plaques, but not soluble Aβ, were associated with impaired CVR (28). More importantly, we find an interaction between elastin haploinsufficiency and AD mutations, such that only the AD mice have impaired hippocampal CVR with elastin haploinsufficiency. These findings suggest that two hits are needed for impaired CVR, a vascular risk factor plus abnormal Aβ or tau.

By influencing neuroinflammation and neurovascular coupling, glia impact CBF. Microglia can play contradictory roles in the brain, what starts as a protective role by removing pathological protein aggregates can transition to a harmful role promoting uncontrolled inflammation(29-31). We found that there were significantly more microglia in the entorhinal cortex of non-AD *Eln*^*+/−*^ mice compared with *Eln*^*+/****+***^ mice and more microglia in the thalamus of AD mice compared with non-AD mice, which is consistent with greater inflammation leading to lower CBF. Astrocytes are primary controllers of neurovascular coupling (32) but can also become reactive and pro-inflammatory(33). As such, our finding of fewer astrocytes in the non-AD *Eln*^*+/−*^ mice is contradictory to our resting CBF and CVR findings, particularly as it has been shown that astrocytes play a role in vasodilation in response to hypercapnia (34). Our finding of considerably more astrocytes in AD mice is in agreement with the current literature (35), and likely represents a heightened inflammatory state rather than a healthy neurovascular environment. In summary, our data potentially suggest that the differences in microglia, rather than astrocytes, have a greater influence on CBF and CVR with elastin haploinsufficiency.

Where we have identified similar effects of elastin haploinsufficiency for males and females, previous reports have indicated that only males are impacted. Hawes et al. examined sex differences in male and female *Eln*^*+/****+***^ and *Eln*^*+/−*^ mice, and found that elastin haploinsufficiency impacted only male mice, specifically for pulse pressure, collagen, and elastin. Similarly, Kailash et al. found that material stiffness was impacted in only male *Eln*^*+/−*^ mice. Conversely, while we find sex differences in CBF and microglia count, these did not interact with *Eln* genotype. These findings potentially suggest that there are sex differences in the effects of elastin haploinsufficiency on large arteries but not the cerebral microvasculature.

Our study is not without limitations. First, the *Eln*^*+/−*^ model is a whole-body genetic manipulation, and thus, we cannot determine if the effects are due to a direct impact of the genotype on the cerebral vasculature or if the cerebrovascular effects are indirectly caused by differences in large artery structure. Second, we did not find evidence of aggregated Aβ or tau in the 3xTg-AD mice. Genetic drift has occurred in this strain over time (36) and a previous study found that male 3xTg-AD mice lack AD-related neuropathology but do have a heightened inflammatory state (37). Thus, our findings in the AD mice may represent an interaction with inflammation rather than an interaction with adverse Aβ and tau. Third, while we studied both male and female AD mice, we did not assess sex differences due to a small sample size. We did not find sex differences for cerebrovascular outcomes in non-AD mice; however, there are sex differences in the rates of AD (38), and thus these are important future analyses. Finally, we did not assess blood CO_2_ to discern if the differences in CVR were due to differences in breathing between the mice.

Overall, our findings suggest that elastin haploinsufficiency leads to lower resting CBF but greater cerebrovascular reactivity, and sex differences do not influence these responses. We further find that elastin haploinsufficiency interacts with AD mutations to impair CVR. These results may suggest that a combination of insults, such as structural vascular changes plus AD genetic risk, are needed for impairment in cerebrovascular function.

## DATA AVAILABILITY

Data will be made available upon reasonable request.

## ACKNOWLEDGMENTS

The 3xTG mice were created by Frank LaFerla at the University of California and obtained from Jackson Laboratory.

## AUTHOR CONTRIBUTIONS

AEW, MMP, and NJA conceived and designed research, AEC, EHR, NRW, GDH, NA, TL, AH, SK, NS, VK, MMP, and RLW performed experiments, AEC, EHR, and AEW analyzed data, AEC, EHR, and AEW interpreted results of experiments, AEC prepared figures, AEC and AEW drafted manuscript, AEC, EHR, NRW, GDH, NA, TL, AH, SK, NS, VK, MMP, NJA, RLW, and AEW edited and revised manuscript and approved final version of manuscript.

